# Hippocampus responds to mismatches with predictions based on episodic memories but not generalised knowledge

**DOI:** 10.1101/2025.02.04.636427

**Authors:** Dominika Varga, Petar Raykov, Elizabeth Jefferies, Aya Ben-Yakov, Chris Bird

## Abstract

Prediction errors drive learning by signalling a mismatch between our expectations and reality. The hippocampus plays a key role in mismatch detection, but it is not known what information the hippocampus uses to form expectations. Here we show that the human hippocampus bases its expectations on episodic memories and not generalised schematic knowledge. Across three fMRI experiments, we demonstrate that the hippocampus is selectively engaged by mismatches with episodic memories of specific events. In contrast, mismatches with schematic knowledge activate regions of the Semantic Control and Multiple Demand Networks, as well as subcortical regions involved in prediction error signalling. Notably, mismatches with episodic memories also engage the Default Mode Network. Overall, our findings provide direct support for some models of learning via mismatch detection and refute models that propose the hippocampus plays a wider role as a more generalised mismatch detector.

## Introduction

Humans possess a remarkable ability to predict what will happen in new situations based on past experiences ^1,2,3^. Detecting a mismatch between our predictions and our in-the-moment experience offers a powerful route to rapidly learn new information (e.g., ^4,5^). The hippocampus plays a central role in mismatch detection (e.g., ^6,7,8,9,10,11^). However, it is not known whether the hippocampus computes mismatches between current experience and predictions made on general schematic knowledge about the past, or episodic memories of specific earlier experiences.

Several proposals suggest that the hippocampus detects novelty by comparing incoming information with stored representations. This includes processing associative mismatch novelty [e.g., where familiar objects are reconfigured into novel arrangements (e.g., ^12^)], contextual novelty [where items are unexpected within a given context (e.g., ^13^)] and schema incongruence [violations of structured knowledge about the world (e.g., ^14^)]. These proposals are based on the well-established preference of the hippocampus for processing relations between items and particularly item-context associations ^15,16,17,18,19^. Furthermore, the neural circuitry within the hippocampus is particularly suited to its hypothesised role as a comparator ^9,20^. However, the prior contextual representations used in this comparator function remain unclear.

There is strong evidence that the hippocampus supports predictions based on specific past experiences to compare to current situations. In humans, it has been shown that the hippocampus detects changes in recently learnt cue-outcomes associations (e.g., ^21,22,23^) and sequences of events (e.g., ^24,25,26^). Likewise, in animal studies, the hippocampus shows a mismatch signal when changes are made to specific, previously encountered environments ^27,28,29^. Some computational models propose that the comparator function of the hippocampus is limited to processing mismatches with episodic-like representations of specific events ^7,30^.

However, often our expectations are based on our generalised understanding of patterns and regularities developed across multiple similar experiences ^31,32,33^. It has been suggested that the hippocampus may also process mismatches based on these more generalised representations ^34,35,36^. This argument is based on evidence that the hippocampus learns the common elements and temporal regularities across multiple past experiences ^37,38,39^, can infer relationships between items that have never been directly experienced together ^40^, and is involved in imagining complex future scenarios ^41^. Some computational models have proposed that the hippocampus plays a key high-level role within a “generative model”, making predictions about the state of the world that are based on abstract generalised knowledge ^10,42^.

The critical test of all these theories of hippocampal mismatch detection is whether the hippocampus responds to prediction errors based on generalised knowledge or whether its role is limited to processing expectations based on episodic-like memories. We address this using three fMRI experiments, where we manipulated the source of expectation violation while participants watched custom-made video clips of actors performing sequences of everyday actions (e.g. doing the laundry). Inside the scanner, all participants watched half of the clips in their ‘Typical’ version (e.g. putting clothes into a washing machine) and the other half in their ‘Atypical’ version (e.g. putting flowers into the washing machine). Depending on participant’s pre-scan familiarity with the clips, actions in the clips mismatched different types of expectations. When participants were unfamiliar with the clips prior to scanning, Atypical actions mismatched solely general Schema Knowledge (Experiment 1). When participants had pre-watched all clips in their Typical version, Atypical actions mismatched both Schema Knowledge and Episodic Memory of the specific clips (Experiment 2). Finally, when participants had pre-watched all clips in their Atypical version, Typical actions mismatched Episodic Memory only (Experiment 3).

Overall, we found strong evidence that the hippocampus is limited to using episodic memory-based representations for its comparator mechanism. Responses to schematic knowledge-based mismatches were found in regions outside of the hippocampus, including cortical control networks and subcortical regions implicated in prediction error processing.

## Results

### The effect of expectation violation on memory

We first analysed memory for target actions, showing high recall accuracy across all experiments (the proportions of remembered target actions are shown in Figure 2a). It is notable that participants’ accuracy significantly increased across the three experiments [β =1.23; 95% CI = [1.09 1.37]; Z =16.85; p < .001; Correct Recall Score ∼ Experiment + (1 | Participant) + (1 | Video Clip)], likely reflecting participants’ increased familiarity with the videos in Experiments 2 and 3 due to the additional pre-scanning phases in these experiments. Interestingly, despite these differences in familiarity, the within experiment logistic mixed-effects models [Correct Recall Score ∼ Condition + (1 | Participant) + (1 | Video Clip)] showed no significant differences in recall accuracy between Typical vs. Atypical actions in any experiment (Experiment 1: β = 0.12; 95% CI = [-0.57 0.81]; Z = 0.33; p = .739; Experiment 2: β = -0.59; 95% CI = [-1.23 0.06]; Z = -1.79; p = .074; Experiment 3: β = 0.42; 95% CI = [-0.31 1.15]; Z = 1.14; p = .254). Overall, participants remembered expected and unexpected target actions equally well, regardless the type of expectation violation.

**Figure 1.**
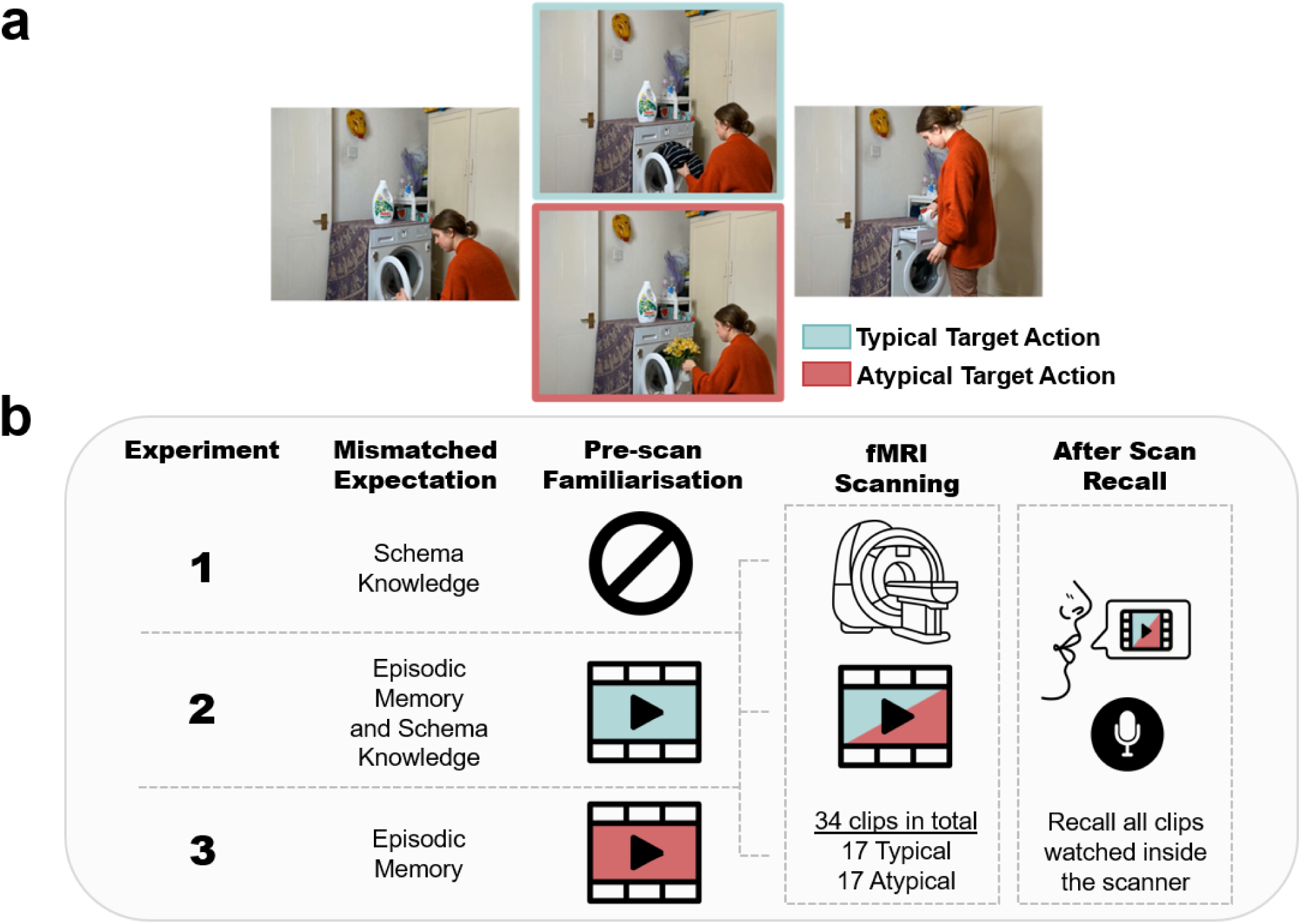
Experimental paradigm. **(a)** Still frames showing example moments from the two alternative versions of the ‘laundry’ video clip (watch the clips <u>here</u>). The two versions showed a nearly identical sequence of actions, except for the target action that was either Typical or Atypical. **(b)** Before scanning, participants either did not watch any clips (Experiment 1), watched the Typical version of each clip (Experiment 2), or watched the Atypical version of each clip (Experiment 3). Across three experiments all participants watched half of the clips in the Typical and the other in the Atypical version during fMRI. By manipulating pre-scan familiarity with the clips, target actions in each experiment violated different types of expectations. After scanning, participants recalled all the video clips watched inside the scanner.

**Figure 2.**
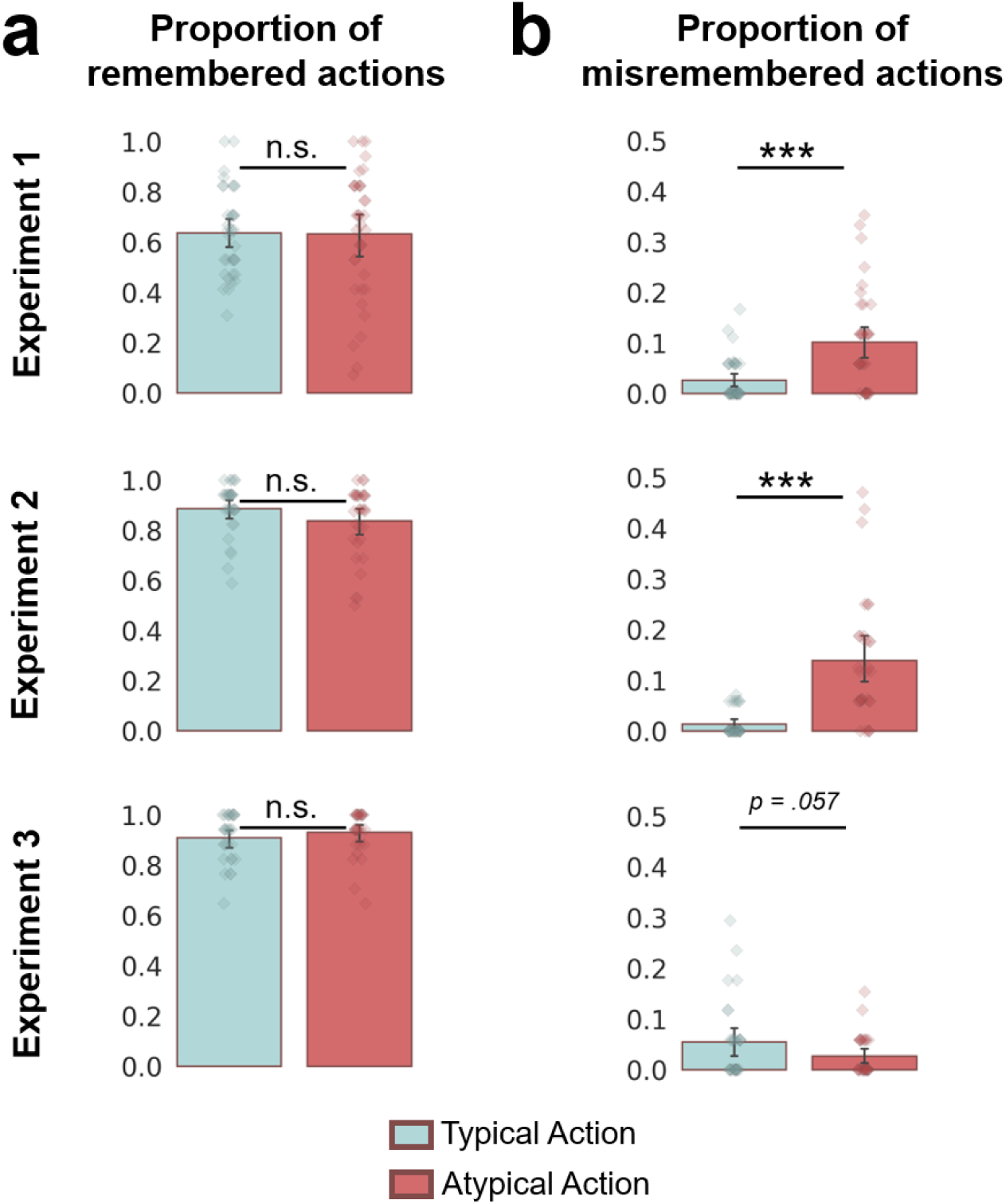
Memory was influenced by the expectedness of actions. **(a)** Bar charts showing the average proportion of remembered target actions in the Typical and Atypical conditions in each experiment. Error Bars represent 95% CI. Strip plots show each participant’s average proportion of remembered target actions for each condition. There were no differences in the average proportion of correctly recalled target actions between conditions. **(b)** Bar charts showing the proportion of erroneously recalled target actions in each condition. People made more errors when recalling actions that violated their expectations. Error Bars represent 95% CI. *** p < .001, n.s. p > .05.

Next, we ran further exploratory analyses to test whether participants were more likely to *misremember* the target actions, depending on whether the action violated expectations (Figure 3b). Participants made considerably more errors when recalling the Atypical compared to the Typical actions in Experiment 1 and 2 (Experiment 1: β = 1.51; 95% CI = [0.77 2.25]; Z = 3.99; p < .001; Experiment 2: β = 2.48; 95% CI = [1.61 3.34]; Z = 5.64; p < .001). There was only a marginally significant difference in the proportion of memory errors between conditions in Experiment 3 [β = -0.81; 95% CI = [1.61 3.34]; Z = -1.91; p = .057; (Recall Error Score ∼ Condition + (1 | Participant) + (1 | Video Clip))]. Interestingly, the effect in Experiment 3 was in the reverse direction to the effects in Experiments 1 and 2; participants were numerically more likely to misremember the Typical actions, having pre-watched versions of the videos showing the Atypical actions. Overall, across all experiments, participants were more likely to misremember unexpected target actions, either by replacing the object involved (e.g., “she put fruits into the washing machine”) or recalling the action without specifying the object (e.g., “she put something strange into the washing machine”).

**Figure 3.**
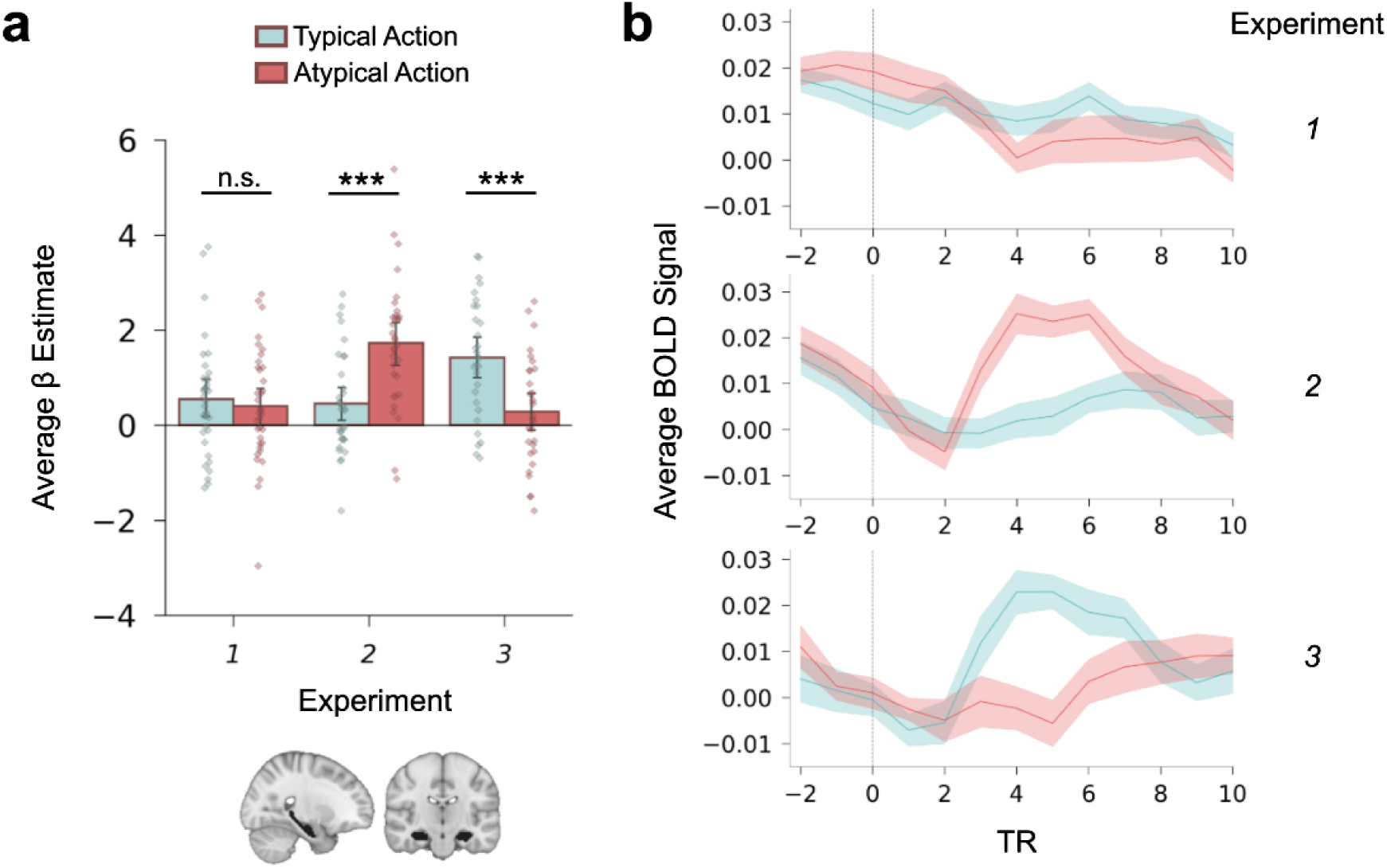
Hippocampal mismatch response is limited to episodic memory-based expectation violation. In Experiment 1, expectations are based only on schematic knowledge, in Experiment 2, expectations are based on both episodic memory and schematic knowledge, in Experiment 3, expectations are based only on episodic memory. **(a)** Bar charts showing the average response to Typical and Atypical target actions in the bilateral hippocampus in the three experiments (strip plots show individual participants’ averaged parameter estimates for the target actions estimated from a GLM). Error bars represent 95% confidence intervals. **(b)** Time course of average BOLD signal in the bilateral hippocampal ROI to Typical and Atypical target actions in each experiment. *** p < .001; n.s. p > .500

In summary, participants remembered the target actions well, and that the actions were remembered differently depending on their expectedness. Importantly, there was no difference in correctly recalling actions that violated or met participants’ expectations. However, participants were more likely to make errors when recalling unexpected actions.

### The effect of different types of expectation violation on hippocampal activity

Our analyses focussed on the hippocampus and the analyses of Experiments 1 (https://osf.io/7g82x) and 3 (https://osf.io/zbnt9) were pre-registered; the analyses of Experiment 2 were not pre-registered but are identical to the other experiments. The analyses aimed to test whether hippocampal activity is modulated by the expectedness of target actions under different types of prior expectations. We used a General Linear Model (GLM) to isolate transient activity evoked by the Typical and Atypical target actions. Hippocampal activity was quantified by averaging beta weights from all voxels within a bilateral hippocampal mask for each condition in each experiment, allowing us to directly compare responses to Atypical and Typical actions. The results are shown in Figure 3.

In Experiment 1, a paired t-test revealed that there was no effect of schema-based expectation violation in the hippocampus (t(35) = 0.57; 95% CI = [-0.36 0.65]; p = .567). Bayesian analysis provided moderate evidence for the null hypothesis (BF_01_=4.78), further suggesting that hippocampal activity was not modulated by target action expectedness in this experiment. To examine whether adding specific episodic memory predictions while watching the same clips would elicit prediction error signals in the hippocampus, we conducted Experiment 2.

In Experiment 2, hippocampal activity showed a transient peak in response to Atypical compared to Typical actions (t(32) = -4.59; 95% CI = [-1.84 -0.71]; p < .001). Bayesian analysis provided decisive evidence for the alternative hypothesis (BF_10_= 378.25), indicating a substantially greater hippocampal response to Atypical actions. This reveals that episodic memory-based predictions are important for eliciting a mismatch response in the hippocampus. However, the increased hippocampal response in Experiment 2 may reflect an additive effect of schema-based and episodic memory mismatches, leaving it unclear whether episodic memory violations alone would elicit increased activity.

To address this concern we conducted Experiment 3, which tested solely episodic memory-based violations. Here, participants pre-watched all the Atypical versions of the videos. Consequently, during scanning, the Typical versions of the videos were unexpected on the basis of episodic memory. In this experiment, hippocampal activity was significantly higher for Typical compared to Atypical actions (t(29) = 5; 95% CI = [0.67 1.60]; p < .001). Bayesian analysis provided decisive evidence for the alternative hypothesis (BF_10_= 910.34), providing compelling evidence that the hippocampus responds to expectation violations even when these violations are driven solely by episodic memory, thus ruling out the influence of schema-based expectations.

Overall, our results demonstrate that the hippocampal role in processing mismatches is limited to expectation violations that are based on episodic memories for specific events.

### The effect of different types of expectation violation outside of the hippocampus

We conducted whole-brain analyses to investigate the mismatch responses outside of the hippocampus in all three experiments. First, we ran separate GLMs for the three experiments, modelling the response to the Atypical and Typical target actions. Group level contrasts between the Atypical and Typical target actions are shown in Figure 4a. We additionally carried out post-hoc ROI analyses investigating whether the effects of expectation violation differed across three brain networks – the Semantic Control Network (SCN), the Multiple Demand Network (MDN), and the Default Mode Network (DMN). We focused on these networks because they are involved in the comprehension of everyday situations and ambiguity resolution ^43,44,45,46,47^. Within each experiment, we conducted a repeated measures ANOVA between the target action conditions and Networks on the average parameter estimates associated with the target actions from all voxels comprising each network (Figure 4b). A final ROI analysis focussed on the ventral tegmental area (VTA) / substantia nigra (SN), since this midbrain region is strongly associated with processing prediction errors and forms part of a functional loop with the hippocampus to process novelty ^9^.

**Figure 4.**
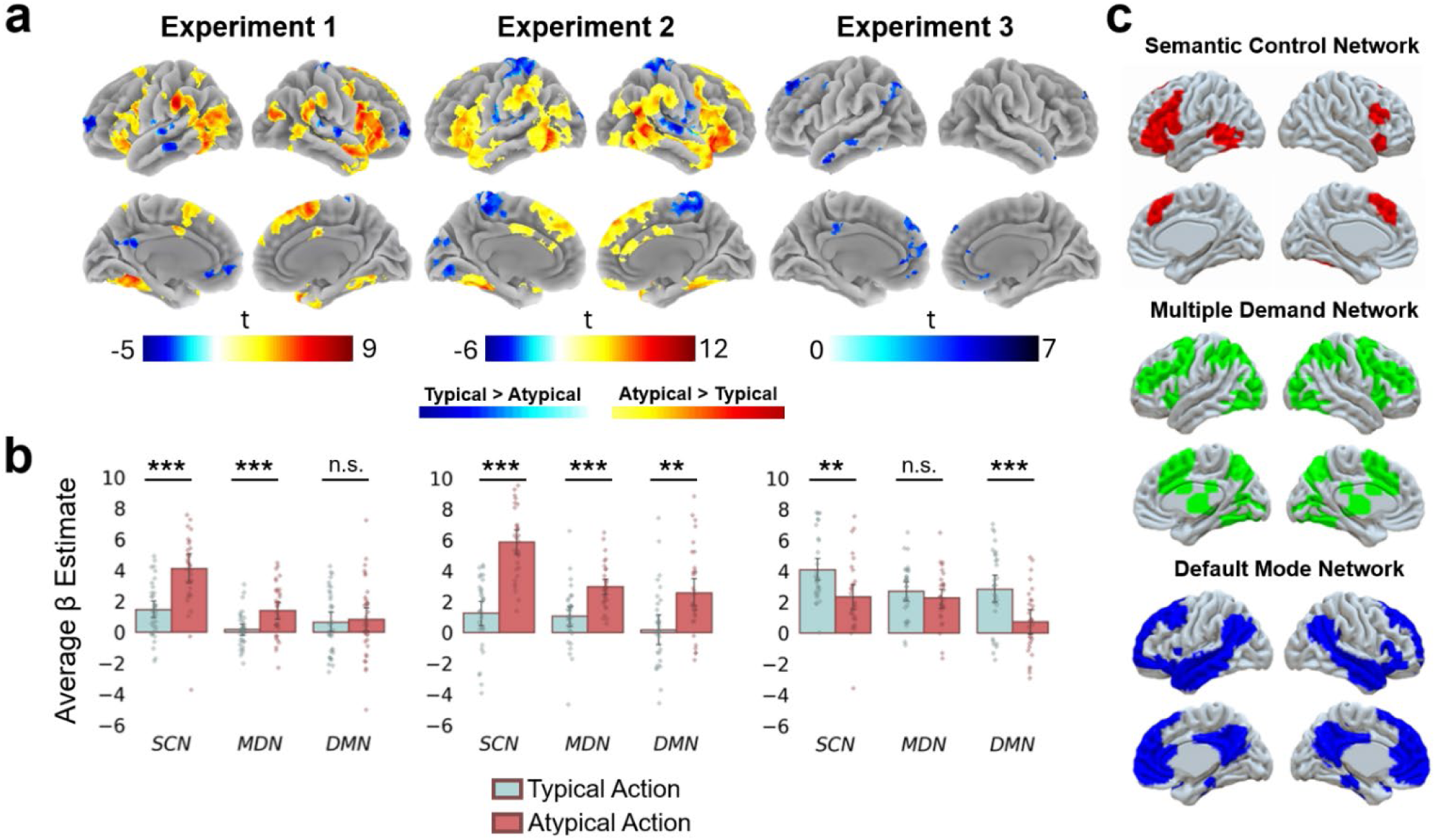
The effect of expectation across the whole brain and in three cortical networks. **(a)** T-maps of the contrast between Atypical and Typical target actions in each experiment. Whole brain t-maps are cluster corrected at FWE p < .05 at voxel height defining threshold of p < .001 and colour-coded to indicate the intensity of activation. The colour bar indicates the t-statistic associated with each voxel. See unthresholded t-maps here: https://neurovault.org/collections/19061/ **(b)** Bar charts show the average response to Typical and Atypical target actions in the Semantic Control Network, Multiple Demand Network and Default Mode Network in the three experiments (strip plots show individual participants’ averaged parameter estimates for the target actions estimated from a GLM). Error bars represent 95% confidence intervals. **(c)** Spatial maps of the network ROIs.

Firstly, in Experiment 1, Atypical compared to Typical actions engaged regions generally implicated in attentional engagement, semantic and predictive processing ^48,49^. Typical compared to Atypical actions engaged regions (such as Posterior Medial and Medial Prefrontal Cortex) implicated in encoding schema-consistent information, comprehending narratives and mentalising ^50,51,52^. This finding is in accordance with suggestions of the SLIMM model (schema-linked interactions between medial prefrontal and medial temporal regions ^14^) that the medial prefrontal cortex has an important role in detecting the match between current events and existing contextual associations. Subcortical effects were also present in the caudate nucleus, which consistent with previous work on expectation violations (particularly with expected actions ^53^), the amygdala ^54^, and the thalamus ^55^.

The network-level analysis revealed significant main effects for Condition (F(1,35) = 13.70; p < .001; eta2[g] = 0.11) and Network (F(2,70) = 35.60; p < .001; eta2[g] = 0.19), and a significant Condition*Network interaction (F(1.65,57.7) = 14.07; p < .001; eta2[g] = 0.06). Pairwise comparisons showed that the average response to Atypical compared to Typical actions was significantly higher in the SCN (t(35) = 5.64; p < .001) and MDN (t(35) = 4.11; p < .001). These networks are implicated in resolving unexpected information by guiding contextually appropriate knowledge retrieval ^56^ and allocating attentional resources to interpret ambiguous situations ^57,58^, respectively. Schema-based expectation violation did not significantly modulate activity in the DMN overall (t(35) = 0.30; p = .763).

Interestingly, there was a significant increase in VTA / SN activity for the Atypical actions, consistent with this region responding to prediction errors ^9,59,60^. A paired t-test comparing Atypical (M=1.08, SD=1.05) and Typical (M=-0.30; SD=1.22) actions revealed an effect of condition, t(35)=2.90, p=.006, 95% CI = [−1.72,−0.30].

In Experiment 2, a strikingly similar map of regions was activated more by Atypical than Typical actions (Figure 4). Once again, subcortical effects were present in the caudate nucleus, thalamus and amygdala. The Network ANOVA revealed significant main effects for Condition (F(1,32) = 57.76; p < .001; eta2[g] = 0.30) and Network (F(2,64) = 24.87; p < .001; eta2[g] = 0.14), and a significant Condition*Network interaction (F(1.29,41.26) = 11.19; p < .001; eta2[g] = 0.06). Pairwise comparisons showed that the average response to Atypical target actions was again significantly higher in regions of the SCN (t(33) = 10.1; p < .001) and MDN (t(33) = 4.95; p < .001). Now with the addition of context-specific memory-based predictions, regions of the DMN were also overall more engaged by Atypical than Typical actions (t(33) = 3.52; p = .001).

In the VTA / SN, there was again greater activity for the Atypical (M=1.08; SD=1.05) compared to Typical (M -0.36; SD=1.22) actions. A paired t-test revealed a highly significant difference, t(32)=5.97, p<.001, 95% CI = [−1.93,−0.95].

The commonality across Atypical actions in Experiment 1 and 2 was that they were surprising with respect to long-term semantic knowledge about the situations depicted in the videos. Therefore, common regions activated in both tasks (in particular regions of the SCN) might reflect violation of predictions based on general knowledge rather than an overarching role in all types of context-violation. Whether these regions would be activated also by context-specific, episodic memory-based violations was tested in Experiment 3.

In Experiment 3, the two sources of expectations – semantic knowledge and episodic memory – are in opposition to each other. Here, unlike the previous experiments, there were no regions more activated by Atypical than Typical actions. This suggests that familiarisation with the Atypical videos rapidly diminished the fMRI response to actions that might be considered inherently surprising based on schematic knowledge. However, several regions that had been engaged by unexpected events in Experiment 1 and 2 (particularly in the dorsomedial and inferior frontal cortex) were now more engaged by Typical than Atypical actions. This reflects the regions responding to events that were unexpected based on memory for the specific videos. The only subcortical structures to also show this effect were the right amygdala and a small region of the right ventral striatum. The Network-related ANOVA revealed significant main effects for Condition (F(1,29) = 19.5; p < .001; eta2[g] = 0.11) and Network (F(1.33,38.59) = 6.87; p = .007; eta2[g] = 0.07), and a significant Condition*Network interaction (F(2,58) = 6.56; p = .003; eta2[g] = 0.03). Pairwise comparisons showed that the average response to Atypical target actions was significantly higher in the SCN (t(29) = 3.56; p = .001), suggesting that these regions likely have a rather general role in processing surprising actions, regardless of the source of expectations. Additionally, the Typical actions also engaged the DMN more than did Atypical actions that matched memory for the specific context (t(29) = 4.72; p < .001). This is consistent with a role for this network when comparing internal representations of recently experienced information with incoming sensory information ^61^. Lastly, expectation violation did not significantly influence activity in the MDN overall in this Experiment 3 (t(29) = 1.29; p = .207). Inspection of the effect in the MDN across all three experiments suggests that in Experiment 3, activity was higher for both the Atypical and Typical actions (compared with Typical actions in Experiments 1 and 2). This is likely due to both types of actions being salient within the clips and capturing participants’ attention.

In the VTA / SN, a paired t-test comparing Typical (M=0.24; SD=1.85) and Atypical (M=-0.21; SD=1.21) actions did not reveal a statistically significant difference, t(29)=1.38, p=.178, 95%CI = [−0.22,1.12].

To summarise, we show that unexpected actions, regardless the source of expectation violation, are processed in regions associated with generating contextually relevant inferences and resolving conflict. At the network level, Atypical, unexpected, target actions that violated general schema-based predictions engaged cognitive control networks more than did Typical, expected, actions. This is consistent with the role of SCN and MDN in processing contextually unexpected information, suggesting that our manipulation of schema-incongruence of the Atypical target actions was successful. Those actions that also violated episodic memory-based expectations, engaged regions of the DMN, which is consistent with the DMN’s role in comparing internally generated inferences (e.g. episodic memories) with incoming information ^51^. Unexpected actions in all experiments were attention grabbing, as suggested by the increased activity in the MDN regions, important for allocating attention to salient, goal-relevant information. Interestingly, subcortical regions associated with prediction error processing were engaged by violations of schema-based expectations, or the combination of schema- and episodic memory-based expectations, but not when expectations were purely based on episodic memory. It is therefore noteworthy that the VTA / SN showed a different pattern of activation across the three experiments compared to the hippocampus.

## Discussion

We performed three separate fMRI experiments to test the role of the hippocampus as a mismatch detector. We found that the hippocampus processes mismatches between expectations based on specific episodic memories and in-the-moment experience. Critically, we show that violations of expectations based on more generalised schematic knowledge do not engage the hippocampus. These findings impose hard constraints on the information used by the hippocampus to detect mismatches. They provide direct support to theoretical models of hippocampal function that identify a more limited role ^7,30^. Conversely, models that have argued for a more general role for the hippocampus in generating and predictions about the state of the world must be reevaluated in the light of the present results (see ^14,36,42^).

Our study showed that Atypical sequences of actions violating long-term schematic knowledge of everyday situations did not differentially engage the hippocampus compared to Typical sequences matching schema knowledge (Experiment 1). Conversely, when participants formed episodic memory-based expectations for the Typical sequences prior to scanning, viewing Atypical sequences in the scanner caused a transient hippocampal response (Experiment 2). This indicates that episodic memory expectations are needed to trigger hippocampal mismatch responses. Nevertheless, it remained a possibility that the additive effect of a mismatch between episodic memory-based expectations *and* general knowledge of the situations shown exceeded a threshold for “surprise” and triggered a hippocampal response. Experiment 3 addressed this possibility, as participants pre-watched all the Atypical versions of the videos before scanning. In this experiment the contextually appropriate Typical versions of the videos caused an increased hippocampal response, since these videos mismatched episodic memories-based expectations from the pre-watch phase. Our findings provide strong empirical support for a family of computational models proposing a role for the hippocampus in comparing incoming information with episodic memories that are stored within the hippocampus ^11,62,63,64^.

Our findings address a key gap in the literature since evidence in support of the hippocampus as a mismatch detector is based overwhelmingly on highly specific, recently learned, information. On the basis of such evidence, the hippocampus has been argued to have a rather general role in processing any information that mismatches expectations based on a context, or even on generalised statistical regularities learnt about the world ^10,13,14,36,37,42^. Our results strongly constrain the role of the hippocampus as a mismatch detector to situations where expectations are based on specific episodic-like representations of the event. Nevertheless, as discussed next, we do not argue that the hippocampus can only make comparisons between the current situation and a memory for an identical experience in the past – indeed such a rigid function would serve little adaptive value in most situations.

It is well-established that the hippocampus is able to support flexible representations that capture the associative structure of a situation or environment and can be used to make inferences about situations that have never been experienced ^65,66,67,68,69^. How do we square the flexibility of hippocampal representations with our results? We suggest that the hippocampus only makes predictions about novel experiences that are part of a learnt "cognitive map”, which may represent a physical space or a more abstract conceptual space ^70^. For example, a map-like representation will encode the relative locations of objects within a particular space, enabling it to detect expectation violations even if exploring the space from an entirely novel perspective ^71,72^. However, when inferences must be drawn from structured mental models abstracted away from a specific cognitive map, mismatches will be detected independent of the hippocampus.

An intriguing question for future research is how closely novel situations need to match prior experiences in order to trigger the hippocampal comparator mechanism. In structured scenarios like visiting restaurants, common features allow us to generalise expectations— such as surveying menus, eating, and paying—but we also must adapt to context-specific variations, like receiving menus via QR codes rather than paper or paying upfront instead of at the end. Abstracting away from the specifics to apply generalised rules to similar situations has been primarily associated with the medial prefrontal cortex ^73^. However, frequently encountered situations that are very predictable and have little variation across similar experiences, such as going through specific security steps in different airports, might allow the hippocampus to leverage these particular contextual associations to generate expectations about novel situations. It remains to be tested how much the underlying structure of different experiences must overlap in order to involve the hippocampus in detecting deviations from expected patterns.

The hippocampal BOLD response in Experiments 2 and 3 might have a number of causes. First, it might reflect the initial increase in activity of hippocampal neurons that signal the mismatch with expectations to other brain regions ^11,28,74^. Alternatively, the detection of a mismatch in the hippocampus might lead to the activation of dopaminergic neurons in the VTA which in turn project back to the hippocampus in order to modulate learning mechanisms within the hippocampus ^9,75^. For example, dopaminergic projections to the hippocampus promote long-term potentiation via activation of D1 receptors ^76^, which in turn has been argued to result in local increases in BOLD activity ^77^. However, this explanation is weakened by the fact that activity in the VTA /SN was driven by violations of schematic knowledge-based expectations, whereas the hippocampus was engaged by episodic memory-based expectation violations. Another possibility is that mismatches trigger release of acetylcholine within the hippocampus which can modulate CA1 synaptic plasticity ^7,78^. Interestingly, acetylcholine can also directly affect the BOLD signal via its action as a vasodilator ^79^. Future research will unpick the cellular mechanisms that underpin the hippocampal mismatch response in our study.

Our results also shed light on the role of various cortical networks in processing unexpected actions. The SCN was strongly engaged when the actions were unexpected based on both schematic knowledge and the combination of schematic knowledge and episodic memory. In Experiment 3, when these sources of expectation were pitched against each other, it was the violation of episodic memories that resulted in significantly higher activity within the SCN. This network plays a role in generating inferences that are appropriate to a particular situational context ^56^. Our findings suggest that the network additionally processes context-specific expectations based on recently experienced information and not only long-term semantic knowledge. This is consistent with previous findings that regions of the SCN control context-appropriate retrieval in both episodic and semantic memory ^80^. The DMN showed a very similar pattern of activation to the hippocampus, suggesting a complementary role in comparing specific episodic memories with current experience. It is possible that hippocampal retrieval of the action watched before scanning triggered reinstatement of its content throughout the DMN (e.g., ^81,82^). We propose that participants may have retrieved the predicted action when viewing an unexpected action, whereas this was unnecessary when observed actions aligned with prior expectations. This is consistent with predictions from a recent computational model, which suggests that episodic memory retrieval preferentially occurs when there is uncertainty about what will happen next ^83^.

Lastly, it is notable that unexpected events were remembered differently to expected events. In Experiments 1 and 2, participants were good at recalling that *something* surprising had happened, but could not necessarily remember what it was (see also ^84^ for similar findings). This revealed itself in an increase in the number of mis-remembered events, which either reflected participants recalling an incorrect action or stating that the action was something strange. Interestingly, correct recall of the target actions was equivalent for both the expected and unexpected videos. This is likely to reflect two opposing effects: surprising actions were inherently more memorable but harder to recall due to their lack of contextual relevance, while typical actions benefited from strong contextual cues (see also ^85^). A previous study used a subset of the same video stimuli as ours, and a recognition paradigm ^86^. Here, the expected and unexpected actions were equivalently cued during the recognition test, and under this situation, the unexpected target actions were remembered better.

In conclusion: our findings demonstrate that the hippocampus computes mismatches between ongoing experiences and stored episodic memories, but not generalised schematic knowledge. The results support theoretical models which have argued for such a limited role for the hippocampus as a comparator, but where direct experimental evidence has been lacking. Conversely, our findings constrain theories that have proposed a wider role for the hippocampus as a more general mismatch detector. Future work must clarify the degree to which hippocampal representations of structured information enable the generation of predictions in similar, yet novel, situations.

## Acknowledgements

This project has received funding from the European Research Council (ERC) under the European Union’s Horizon 2020 research and innovation programme, grant number: 819526 to C. M. B., and also from Sussex Neuroscience, and from a University of Sussex, School of Psychology PhD scholarship to D.V. We thank Sam Berens and other members of the Sussex Episodic Memory Group, as well as Neil Burgess, Caswell Barry, Mick Rugg and John Aggleton for helpful comments and discussions. We thank colleagues at the Clinical Imaging Sciences Centre for their support in fMRI scanning.

## Author Contributions

D. V. and C. M. B. conceived the experiment. A. B. Y. contributed to the experimental design early on. D. V. programmed the experiment, collected all data, scored the recall data, and performed the analyses. P.R. provided support for the analyses, P. R., A. B. Y. and E. J. provided analysis ideas. D.V. and C. M. B. wrote the original manuscript. All authors discussed the results and edited the manuscript.

## Declaration of Interests

The other authors declare no competing interests.

## Resource Availability

### Lead Contact

Further information and requests for resources should be directed to and will be fulfilled by the Lead Contact, Dominika Varga.

### Materials Availability

The video clips are available upon publication on the Open Science Framework. Please e-mail the Lead Contact for other information about the materials.

### Pre-registration

For Experiment 1 and Experiment 3, the rationale, hypotheses, study design and detailed analysis plan were pre-registered (Experiment 1 Pre-registration link: https://osf.io/7g82x; Experiment 3 Pre-registration link https://osf.io/zbnt9). Experiment 2 was not pre-registered, but uses the same materials, has a similar study design and uses identical fMRI analyses as Experiment 1 and 3.

### Data and Code Availability

All code and data from this project will be openly available upon publication on the Open Science Framework project link: https://osf.io/p6z2g. Group level contrasts between Atypical and Typical actions are on neurovault.org: https://identifiers.org/neurovault.collection:19061.

## Methods

### Participants

There were three separate groups of healthy adult participants recruited for the three experiments. An a priori power analysis for the hippocampal contrast of Atypical vs Typical actions, suggested that 30 participants were required to detect a Cohen’s d effect size of 0.4, at a level of α = 0.05 and with a power of 0.7. A Cohen’s d effect size of 0.4 is consistent with the effect of the modulation of hippocampal response to surprising events by prediction strength reported by Long, Lee & Kuhl (^23^). For Experiment 1 thirty-seven participants were recruited (29 female, 8 male, range = 18-30 years old). One participant was excluded from any further fMRI data analysis due to issues with MRI data acquisition. In total, there were thirty-six participants included in the final dataset in Experiment 1 (29 female, 7 male, mean age = 21.6 years old, SD = 3.3 years). For Experiment 2 thirty-seven participants were recruited (25 female, 12 male, range = 18-32 years old). Two participants were excluded from further fMRI analysis due to poor MRI data quality (excess signal dropout), and two due to not following the instructions. In total there were thirty-three participants included in the final dataset in Experiment 2 (22 female, 11 male, mean age = 22 years old, SD = 3.4 years). For Experiment 3 thirty participants were recruited (25 female, 5 male, mean age = 21 years old, SD = 2.8 years, range = 18-29 years old). All participants had normal or corrected-to-normal vision, were right-handed, fluent English speakers. No participants were taking prescribed medication for a mental health condition. Participants were recruited via campus flyers and advertisements posted on the School of Psychology’s online participant recruitment system (Sona Systems). Informed consent was provided by all participants before the experiment, and they were given monetary compensation for participating (£10/hr). All three experiments were approved by the Brighton and Sussex Medical School Research Governance and Ethics Committee.

### Stimuli

Custom-made stop-motion video clips were used in all experiments. Each video clip showed an individual person undertaking a series of actions in an everyday situation (such as doing the laundry). A range of actors were used throughout the different videos. The clips took place in different locations and depicted different scenarios. The camera maintained a fixed viewpoint throughout each scene. Thirty-four scenarios were used, and each scenario had two alternative versions resulting in 34 pairs of clips in total. 10 pairs were the same as those used in a previous study by Ben-Yakov and colleagues (^86^) and are available at https://osf.io/rjav4. The other 24 pairs were created to be similar to the original clips in terms of length and format. The clips within each pair were identical to each other, except for one action (which we will refer to as the target Action). In one version, the target action was Typical, fitting with the overall situation, while in the other version the target action was Atypical, incongruent with the context. For example, in the Typical condition of the laundry video the actor was loading a washing machine with clothes, while in the Atypical condition the actor was loading a washing machine with flowers. For each pair of clips, the timing of each action was identical, and apart from the target action, every other action was the same (e.g., entering the room, putting the washing liquid on top of the washing machine, opening the drum of the machine, closing the door after loading the machine, putting the washing liquid into the machine’s drawer, etc.). For Experiment 1, which investigated surprise based on Schematic Knowledge, each video clip included two scenes: one scene with a target action (the Target Scene, average length=30 seconds) and another scene with no target action (the Non-Target Scene, average length=17 seconds). The Non-Target Scene was always related to the Target Scene but set in a different location and the person was carrying out solely context appropriate actions (such as folding clothes in a bedroom). The order of Target and Non-Target scenes were counterbalanced across the 34 clips (17 clips with Target Scene presented before, and 17 clips with the Target Scene presented after the Non-Target Scene). The reason for including Non-Target scenes in the original study of Ben-Yakov et al., (^86^) was to test whether surprising target actions affected memory for actions within the Non-Target scene (which it did not). We included the Non-Target scenes in Experiment 1 in order to investigate BOLD activity changes at scene changes. However, we do not report these results here. For Experiment 2 and 3, each video clip comprised only the Target Scene.

### Procedure

Across the three experiments we manipulated the typicality of sequences of actions shown within the video clips inside the scanner as well as participants’ familiarity with the video clips prior to scanning.

#### Pre-scanning Session

##### Experiment 1

Before entering the fMRI scanner, participants in the Schematic Knowledge based mismatch group were only briefed about the structure of the task that they will carry out inside the scanner by watching a PowerPoint presentation that explained the format of the scanner task. Importantly, participants in Experiment 1 did not watch any video clips before entering the scanner.

##### Experiment 2

Before entering the fMRI scanner, participants were familiarised with the Typical version of each video clip. First, participants watched the clips in a randomised order. Second, they were asked to recall each clip in as much detail as possible, especially focusing on the sequence of actions carried out by the actors. Participants were cued with an image of the first frame of each clip and asked to say out loud what they remembered happening in the clips. This recall phase was self-paced with the opportunity to take a short break between recalling each clip. The order of clips in the recall phase was randomised across participants. Third, after recalling each clip, they watched each video clip once again to refresh their memory. The pre-scanning session took 66 minutes to complete on average. At the end of this video familiarisation phase, participants were told that inside the scanner they will see some clips identical to the ones they had just watched and some clips that will contain a change.

##### Experiment 3

The pre-scanning procedure in Experiment 3 was similar to Experiment 2, except that participants watched all clips in their Atypical version, showing the Atypical target action in each clip. The memory training had identical structure to Experiment 2. The pre-scanning session took 66 minutes to complete on average.

#### Scanning Session

Inside the scanner, participants in all three experiments watched 34 unique clips in total – half of the clips in the Typical condition and half in the Atypical condition. Due to an issue with the script that ran the stimuli in Experiment 2, 15 (out of 33) participants watched 18 clips in the Typical version and 16 in the Atypical version of the clips. Across participants we counterbalanced which videos were seen in the Atypical and which clips were seen in the Typical condition. Each clip was seen only once during the scanning session and presentation order was randomised. In all three experiments the scanning session lasted about an hour. The scanning task was carried out across 5 functional runs (Experiment 1 average length of runs = 8 minutes, Experiment 2 and Experiment 3 average length of runs = 6 minutes). Four runs contained 7 trials and 1 run contained 6 trials. A trial refers to a sequence of fixation-video-fixation-question--fixation-odd/even task. Each trial started with a red fixation cross that signalled the upcoming video (Experiment 1: Average Length = 1.30s, SD = ±0.06s; Experiment 2: Average Length = 1.29s, SD = ±0.07s; Experiment 3: Average Length = 1.35s, SD = ±0.15s). After the red fixation cross, each video started at the start of a TR. After watching a clip, participants saw a brief fixation cross (Experiment 1: Average Length = 2.22s, SD = ±0.10s; Experiment 2: Average Length = 2.22s, SD = ±0.08s; Experiment 3: Average Length = 2.19s, SD = ±0.08s) and were then asked a short Yes/No question related to the clip they just watched, which was presented for 5 seconds maximum (Experiment 1: Average RT = 2.03s, SD = ±0.37s; Experiment 2: Average RT = 0.78s, SD = ±0.22s; Experiment 3: Average RT = 0.88s, SD = ±0.24s). This was followed by a brief fixation cross again (1.5s for all trials in all experiments). Each trial ended with an Odd/Even number judging task (for 10s in all experiments). Responses to the question and the number judging task were made using a two-button response box held in the left hand (“Yes” and “Odd” with their middle finger, "No” and “Even” with their index finger). This overall task structure was consistent across all three experiments.

Participants in Experiment 1 were instructed to pay attention to the clips as they would be asked to recall them after the scanning session. The in-scanner questions assessed comprehension of the preceding video clip (e.g. half of the participants were asked “Did the actor use a washing machine?” [correct answer is “Yes”], while the other half were asked “Did the actor use a dishwasher?” [correct answer is “No”]). Participants in Experiment 2 and 3 were instructed to pay attention to the clips as they would be asked if they noticed any change to the clips compared to what they had seen prior to scanning and that they will have to recall the clips shown in the scanner after scanning. In Experiment 2, video clips in the Typical condition were identical to those watched before scanning, but video clips in the Atypical condition replaced the previously seen Typical target action with an Atypical target action. Conversely, in Experiment 3, video clips in the Atypical condition were identical to those watched before scanning, but video clips in the Typical condition replaced the previously seen Atypical target action with a Typical target action. At the question phase, participants were asked, “Have you noticed any change in this clip?” and made a “Yes”/“No” response via the button box.

#### Post-scanning Session

In all three experiments, participants were asked after scanning to recall what happened in the video clips they watched inside the scanner. Their recall was audio recorded. Participants were cued with a picture of the first frame in the Target scenes and were asked to recall out loud what happened in the video clip, particularly focusing on the sequence of actions the actor was performing. The recall task was self-paced, participants were given the opportunity to take a short break between recalling each clip. Note that in Experiment 1, the Target scenes that were cued were half of the time presented as the second scene in the video clips inside the scanner. Therefore, for half of the clips the cue did not indicate the beginning of the whole video clip, but the middle, where the scene changed. This was because our primary aim was to measure recall of the target actions; therefore, we aimed to optimise recall performance for the part of the clips that showed these actions.

### fMRI Acquisition and Pre-processing

All images were acquired on a 3-T Siemens Prisma scanner with a 32-channel head coil. To minimize head movement, soft cushions were inserted into the head coil. Functional images were acquired with a gradient-echo EPI sequence with multiband acceleration factor of 3 with the following parameters (TR 1520 ms, TE 28 ms, 75° flip angle, field of view = 208mm x 208mm, 72 slices with slice thickness of 2 mm and isotropic 2 mm voxels). Two SpinEcho fieldmap runs with reversed phase-encode blips in both anterior to posterior and posterior to anterior were acquired with the same parameters as the functional images. Separate field maps were acquired for each functional run. A high -resolution T1-weighted image was acquired with 3-D MPRAGE sequence (in Experiment 1 and 2: TR 2530 ms, TE 1.63 ms, 7° flip angle, field of view = 240mm x 256mm with slice thickness of 1mm and 1 mm isotropic voxels; in Experiment 3: TR 2300 ms, TE 2.19 ms, 9° flip angle, field of view = 256mm x 256mm with slice thickness of 1mm and 1 mm isotropic voxels). Preprocessing of structural and functional data were carried out using standard processing pipeline (fMRIPrep) that was developed by Esteban and colleagues to increase robustness and replicability of fMRI results ^87^. Experiment 1 used fMRIPrep version 20.2.1, Experiment 2 and Experiment 3 used fMRIPrep version 21.0.0. In short, pre-processing steps involved skull-stripping the T1-weighted (T1w) image, and segmenting it into cerebrospinal fluid (CSF), white--matter (WM) and gray-matter (GM). Then the brain-extracted T1w image was registered to an MNI152 template. The BOLD functional runs were skull -stripped, head-motion parameters were estimated over the series, and slice-time correction was applied. Each BOLD timeseries was then co-registered to the native T1w image. Several confounding time-series were calculated, including framewise displacement (FD), the derivative of variance (DVARS) over frame--to-frame motion, and three region-wise global signals (from CSF, WM, and whole-brain masks). Lastly, the BOLD time-series were resampled into standard MNI152NLin2009cAsym space. Pre-processed MRI data were further analysed using MATLAB (2020b) and Statistical Parametric Mapping software (SPM12; Wellcome Trust Centre for Neuroimaging, London, UK). We included six motion parameters, framewise displacement, White Matter Signal and Cerebrospinal Fluid Signal as regressors of no interest in all first level models to account for any residual noise and motion effects after motion realignment. For whole-brain analysis, functional data were smoothed with a 6 mm FWHM kernel. For hippocampal and VTA/SN Region of Interest analyses, functional data were smoothed with a 3 mm FWHM kernel.

### Analyses

#### Behavioural Data Analysis

Behavioural data were collected after scanning of participants’ recall of all the video clips they watched inside the scanner. Recall scoring was identical for all experiments.

#### Recall Scoring

In Experiment 1, recall data from 36 participants were analysed. In Experiment 2, data from 31 participants were analysed (1 excluded due to problems at the scanning stage, 2 excluded due to not following instructions, 3 lost data). In Experiment 3, data from 30 participants were analysed. Responses were scored by one of the authors (Dominika Varga). Data scoring focused only on the target actions; other recalled details about the video were not scored. Each target action was made up of one meaningful piece of action (for example, brushing teeth) carried out with either an expected object (e.g., toothbrush) or an unexpected object (e.g., rhubarb).

##### Remembered / Forgotten Scoring

Target actions were first scored as remembered (score 1) or forgotten (score 0). A target action was scored “remembered” if the participant described the action itself and the object it was carried out with. An exception to this were Typical target actions that could realistically only be performed using a specific object (such as brushing teeth with a toothbrush). Here, responses were scored as remembered even if the object was not mentioned (e.g. “the actor brushed their teeth”).

##### Memory Error Scoring

We further investigated whether trials that were scored “forgotten” were simply due to the participant omitting the target action entirely from their recall or having an imperfect recall of the target action. We scored two types of memory errors associated with the imperfect recall of a target action, an “Unspecified” Target memory error and a “Replaced” Target memory error. A trial was scored Unspecified Target either if the action was mentioned but the object was left out or replaced with the word “something”, or, in the case of the Atypical Targets specifically, if the participant stated that the object was weird/unexpected but could not recall its specific identity. A trial was scored Replaced Target if the participant recalled the correct action but replaced the object with another object. The remainder of the forgotten trials were classified as omissions (the target action was not mentioned during recall) – i.e. were not counted as errors.

#### Recall Analysis

Data from all three experiments were analysed using logistic mixed effect models estimated with the lme4 ^88^ package available in R. First, to compare difference in memory across typical and atypical conditions, separate models were fitted for each experiment. Then to compare overall memory accuracy across the three experiments, data were combined across all experiments. Figures were created using Matplotlib and Seaborn libraries in Python.

Note that in Experiment 1 there were 10 participants who had one or more missing recall trials due to issues with the recording equipment or due to not having watched all the videos in the fMRI scanning session (percentage of trials missing across all 36 participants: M = 10%, SD = ± 23%). In Experiment 2 the 15 participants with imbalanced conditions had one target action incorrectly presented in the Typical Condition (see Scanning Session under Procedure). This incorrect trial was coded as a missing trial. 1 other participant had six trials coded as missing because they did not watch these clips inside the scanner. Missing trials were excluded from the logistic mixed effects models. There were no missing trials in Experiment 3.

The following analyses was identical across all three experiments.

##### Remembered/Forgotten Target Action Analysis

Our main interest was to test whether there is a difference in recalling the target actions correctly depending on whether the target was in the Typical or Atypical condition. Therefore, we gave a binary memory score of remembered (1) or forgotten (0) for each trial.

To examine whether the Typical and Atypical conditions affected the proportion of remembered targets, we ran a mixed-effect logistic regression. We entered the binary memory score as the dependent variable and included a predictor indexing whether the target action was in the Typical (0) or Atypical (1) condition and random intercepts for participants and video clips [Correct Recall Score ∼ Condition + (1 | Participant) + (1 | Video Clip)].

To compare overall memory accuracy across experiments, we entered the binary memory score as the dependent variable and included a predictor indexing the Experiment (1,2,3) and random intercepts for participants and video clips [Correct Recall Score ∼ Experiment + (1 | Participant) + (1 | Video Clip)]. This analysis was not preregistered.

##### Memory Errors Analysis

We wanted to see whether the experimental condition affected the type of errors people made. Specifically, we tested whether participants were more likely to recall something about the target actions that was incorrect depending on whether the action was Typical or Atypical. A score of 1 was entered into the analysis for those forgotten trials coded as either Replaced target actions or Unspecified target actions. The rest of forgotten trials and remembered trials were entered into the analysis with a score of 0 (no error). The trials that were missing were excluded.

To examine whether the Typical and Atypical conditions affected the proportion targets erroneously recalled, we ran a mixed-effect logistic regression in each experiment. We entered the binary error score as the dependent variable and included a predictor indexing whether the target action was in the Typical (0) or Atypical (1) condition and random intercepts for participants and video clips [Memory Error Score ∼ Condition + (1 | Participant) + (1 | Video Clip)]. This analysis was not pre-registered.

### fMRI Data Analysis

#### General Linear Model

##### Experiment 1 General Linear Model

The design matrix of the General Linear Model (GLM) included five regressors. The focus of our main analyses was on transient changes in activity evoked by key moments in the task. For this reason, we included regressors for: 1) the onset of the video clips; 2) the scene changes; 3) Atypical target actions; 4) Typical target actions; 5) the full duration of the comprehension questions and following fixation cross. The first four regressors had zero duration (delta functions). The onset of each target action (Atypical or Typical) corresponded to the most surprising timepoint of the Atypical version of each pair of clips. To determine the most surprising timepoints, we asked 18 independent raters to indicate the moment they found most surprising while watching the Atypical versions of each video clip. For each clip, responses that were given within the target action’s duration were averaged together to give the most surprising timepoint of that clip. These timepoints were then used as the onset for the corresponding Atypical and Typical versions of target actions in our analysis. On average, these most surprising timepoints were 1.4 seconds after the actual onset of the target actions. The odd/even number judging task, the rest of the movie time-points and the fixation cross at the beginning of each trial were unmodeled, acting as the baseline.

##### Experiment 2 & 3 General Linear Model

The design matrix of the GLM included four regressors. The regressors included the 1) video clips onsets, 2) Atypical and 3) Typical target actions, and the 4) comprehension question with the inter-trial fixation crosses. All four regressors were modelled identically to Experiment 1. The only difference between the two GLMs was the absence of a scene change regressor in Experiment 2 and 3 (hence the absence of scene changes in the video clips in Experiment 2 and 3). The odd/even number judging task, the rest of the movie timepoints and the fixation cross at the beginning of each trial were unmodeled, acting as the baseline.

#### Statistical Thresholding

For whole brain analyses, group level testing was done using a one-sample t-test on the functional maps generated by the first level analysis. Whole brain maps were cluster corrected at FWE p < .05 at voxel height defining threshold of p < .001.

#### Region of Interest Analyses

In all ROI analyses, after estimating the first level whole brain GLMs for each participant (see GLM above) we used FSLUTILS’ fslmeants program to average the beta weights associated with the Atypical and Typical target actions from all voxels within the ROI.

##### Hippocampal ROI

The hippocampal ROIs were taken from https://neurovault.org/collections/3731/. The main analyses included the combination of left and right “head”, “body” and “tail” ROIs into one unified bilateral hippocampal ROI.

For each experiment we performed planned two-tailed paired t-test comparing BOLD differences between Typical and Atypical actions. We also report complimentary Bayes Factors using two-tailed standard Cauchy prior with scale 0.707.

##### Network ROIs

We used two control networks defined from previous studies: the semantic control network (SCN) ^89^ and multiple-demand network (MDN) ^58^. Maps of these networks were decomposed into two non-overlapping maps containing mutually exclusive SCN regions and MDN regions ^90^. A map of Default Mode Network (DMN) was taken from the 7-network parcellation from Yeo et al. (^91^).

To compare BOLD response to target actions across the Network ROIs, we conducted a repeated measures ANOVA within each experiment, with within subject factors: Condition (Typical and Atypical target actions) and Network (Semantic Control, Multiple Demand, Default Mode Network). ANOVAs were conducted with the ez package available in R ^92^. In case the ANOVAs showed significant results, post-hoc two-tailed paired t-tests were conducted between all sessions/groups, with Bonferroni correction applied to account for multiple comparisons.

##### Ventral tegmental Area (VTA) / substantia nigra (SN) ROI

The VTA / SN ROI was taken from https://neurovault.org/images/786456/. For each experiment we performed two-tailed paired t-test comparing BOLD differences between Typical and Atypical actions.

#### Activity Time Course Analysis

We first regressed out variation due to head motion with 6 Motion Parameters, Framewise Displacement, White Matter and Cerebro Spinal Fluid signal from each functional run. Then from these filtered functional runs, we extracted and normalized (z-score) the time course from the bilateral hippocampal ROI. For each trial, we interpolated the time course to align with the true onset of the target actions. The resolution of the time course was kept as the TR resolution of 1.52 seconds. We then binned the time course around each target action TR by TR (in steps of 1.52 seconds) starting from 2 TRs before the onset of the target action and ending 10 TRs after the onset of the target action (in total 13 TRs time course window for each trial). For plotting, we averaged across each time bin for each condition within participants to get an average BOLD Signal time course for the Typical and the Atypical target actions.

## Notes

### Competing Interest Statement

The authors have declared no competing interest.

